# Lung surfactants as a component of lipid nanoparticles for pulmonary mRNA delivery

**DOI:** 10.64898/2026.05.20.726526

**Authors:** Sarah S. Nasr, Owen E. Tabah, Sahana Kumar, Gregg A. Duncan

## Abstract

Pulmonary delivery of lipid nanoparticles (LNPs) remains an area of significant interest, given the broad range of genetic disorders that could be addressed through localized administration of therapeutic nucleic acids to the lung. In this study, we investigated how incorporation of the clinically used lung surfactant cocktail Poractant alfa affects the *in vitro* and *in vivo* transfection performance of mRNA-loaded LNPs. The resulting lung surfactant–enhanced LNPs (Surf-LNPs) exhibited substantial improvements in particle assembly, yielding an order of magnitude higher particle concentration at equivalent input conditions compared to conventional (Onpattro-like) LNP formulations. In vitro, Surf-LNPs demonstrated several-fold increases in mRNA transfection efficiency and protein expression while maintaining excellent cytocompatibility. These enhancements are attributed to an elevated apparent pKa and the surface-active properties of surfactant protein B (SP-B), which promote more rapid and efficient endosomal escape relative to conventional LNPs. *In vivo* evaluation following intranasal administration further revealed enhanced mCherry expression in the lungs of mice treated with Surf-LNPs compared to conventional LNPs. Ultimately, these findings establish lung surfactant incorporation as a simple yet powerful formulation strategy to improve pulmonary gene delivery using LNPs, with the potential to significantly advance the translation of inhaled nucleic acid therapeutics.

---

Lipid nanoparticles (LNPs) have emerged as a first-in-class pharmaceutical agent for nucleic acid delivery. Cationic and ionizable lipids have been widely explored in LNP formulations for gene delivery, with the latter being the most widely used for therapeutic mRNA delivery^1^. An important feature of LNPs is their tunability, depending on the chemical structures of the ionizable lipid and other lipid components, which can alter their uptake in specific cell types and their trafficking to specific organs^2–6^. Despite these desirable characteristics, pulmonary delivery of LNPs has been met with challenges due to their inflammatory potential when delivered via intranasal and inhaled routes ^7–9^. To address these challenges, several groups have engineered new classes of ionizable lipids with properties that promote transfection within the lung epithelium to reduce dose requirements to achieve a therapeutic effect^10–12^. To formulate LNPs capable of resisting shear-induced disruption following nebulization^13^, modifications to LNP formulations can be made, including but not limited to increasing the molar amount of PEGylated lipid^14,15^, optimization of LNP assembly or dialysis buffers^16^, and inclusion of other stabilizing excipients such as alcohols, sugars, peptides, nonionic surfactants, or supramolecular DNA hydrogels ^16–20^. Beyond the traditional LNP components, other additives, such as cell-penetrating peptides have been incorporated to enhance delivery to target cells via the pulmonary route ^21^. This body of work illustrates the complexity of generating LNPs with the appropriate attributes to be administered via the pulmonary route, ability to overcome biological barriers to delivery such as airway mucus, and capacity to deliver mRNA cargo to disease-relevant cell types in the lung. Additionally, improving LNP performance in cellular uptake and endosomal escape can help offset the reduced efficiency in these functions caused by the high level of PEGylation required for effective penetration through the mucus barrier^22–24^.

In this context, lung surfactants produced by type II alveolar epithelial cells have been dually considered as agents to enhance pulmonary nanoparticle delivery^25–27^, and conversely, as a barrier to effective nanoparticle delivery to the respiratory tract^28^. For example, prior work has shown lung surfactant coatings on NPs can alter their biodistribution and reduce their uptake in phagocytes such as alveolar macrophages^25^. We hypothesized, based on the structural similarities between lung surfactants and components of conventional LNPs, specifically, helper lipids and cholesterol, that efficacious lung surfactant-based LNPs can be formulated that are better suited for pulmonary gene therapy applications. To test this, we engineered LNPs where distearoylphosphatidylcholine (DSPC), the standard helper lipid in many commercially available LNPs, was replaced with Poractant-α, a porcine lung surfactant cocktail (**Table 1; Fig 1A**). We will refer to these Poractant-α containing LNPs as “Surf-LNPs” throughout the manuscript. Poractant-α is a natural lung surfactant used to treat respiratory distress syndrome (RDS) in premature infants^29^ and contains a physiologically relevant assortment of phospholipids and lipoproteins **(Fig S1-A)**^30^. Dipalmitoyl phosphatidylcholine (DPPC) is the main phospholipid component in poractant-α (37.5% molar ratio) and is very similar to DSPC. DSPC and DPPC are both saturated phospholipids with the same choline headgroup, with the only structural difference being a shorter (C_16_) hydrocarbon tail in the case of DPPC compared to DSPC (C_18_) **(Fig S1-B)**.

**Table 1.** Composition of LNPs and Surf-LNPs, represented as % molar ratios of the enlisted lipids and the universal N/P ratio used to prepare both particle types where: Dlin-MC3-DMA = Dilinoleyl-methyl-4-dimethylaminobutyrate; DSPE=distearoyl-sn-glycero-3-phosphoethanolamine; DSPC = distearoylphosphatidylcholine; PEG2000: polyethylene glycol (2 kDa MW); N/P ratio = Nitrogen-to-phosphate molar ratio.

|  | LNP | Surf-LNP |
| --- | --- | --- |
| Dlin-MC3-DMA | 50% | 50% |
| Cholesterol | 38.5% | 38.5% |
| DSPE-PEG2000K | 1.5% | 1.5% |
| DSPC | 10% | — |
| Poractant-α | — | 10% (calculated as DPPC) |
| N/P ratio | 6 | 6 |

**Figure 1.**
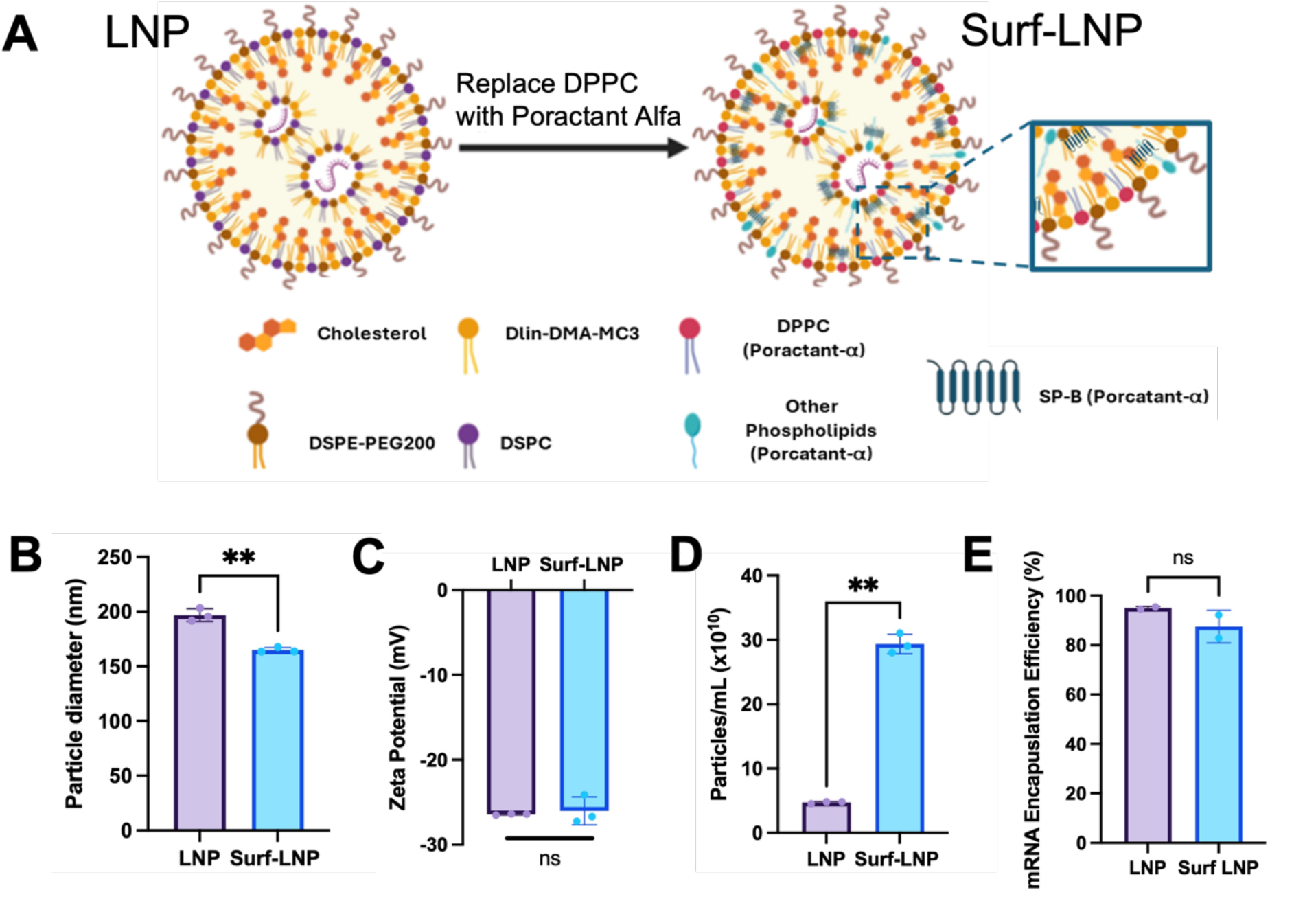
Physicochemical characterization of lung surfactant enhanced lipid nanoparticles (Surf-LNP). (**A**) Diagrammatic illustration of LNP vs Surf-LNP composition. Physicochemical properties of Surf-LNPs vs LNPs Characterized using Nanoparticle Tracking Analysis (NTA) (**B**) Particle diameter (nm). (**C**) Zeta potential (mV), (**D**) Particle count (particles/mL). (**E**) mEGFP encapsulation efficiency in LNP vs Surf-LNPs assessed using Ribogreen assay. Data sets statistically compared by student’s t Test: **p < 0.01; ns = not significant.

We would also like to credit prior work by Merckx et al.^31^ and Guagliardo et al.^32^ which further inspired the approach taken for the studies herein. Merckx et al. used either a lipid cocktail or poractant-α to coat an siRNA-dextran nanogel. The lipid cocktail used was DPPC and PG, the main phospholipid components of porcine surfactant alpha, making SP-B and SP-C the only unique components when compared to poractant-α. Yet they showed that, while the lipid cocktail dramatically reduced particle uptake and gene silencing activity compared to non-coated particles *in vitro*, poractant-α reduced uptake to a slightly lesser extent and maintained gene silencing efficiency that was almost equivalent to that of the non-coated particles. Similarly, *in vitro* results were also observed when gelatin-DNA nanogels were coated with an LNP lipid mixture; the coated particles showed negligible transfection efficiency^33^. The gene silencing capacity *in vivo* for this nanogel system was most pronounced for SP-B-containing lipid cocktails, which were closer in composition to the full poractant-α cocktail, whereas transfection by non-coated nanogels was virtually undetectable. Nanogels coated with the lipid cocktail exhibited only a modest gene-silencing effect compared to their SP-B-containing counterparts. They thus demonstrated that SP-B was an efficient RNA delivery enhancer when incorporated into a suitable phospholipid mixture.

Surfactant Protein B (SP-B) is a strongly hydrophobic protein, yet it also contains a substantial fraction of basic amino acids, lending it a net positive charge. Its mature form is a homodimer featuring two chains, each 79 amino acids long^34^. Along with the lipid components of lung surfactant, SP-B plays a crucial role in stabilizing the surface tension at the pulmonary air-liquid interface, preventing lung collapse during respiration^35,36^. Each monomer of the homodimer is comprised of 4–5 amphipathic α-helices, stabilized by three intramolecular disulfide bonds. The dimer is then formed by one intermolecular disulfide bond between the two monomers^37^. This tertiary structure is homologous to saposin-like proteins (SALIPs)^38^. The characteristic SALIP fold comprises 4-5 α-helical hairpin units that facilitate its interaction with lipid membranes^39^. Where the hydrophobic axes of the helices featuring mainly the hydrophobic amino acids are oriented parallel to the lipid tail’s interface, and the protein’s positive charges, endowed by the basic amino acids, are thought to interact with the anionic heads of phospholipids^34,40,41^. We hypothesized that this could represent the primary mode of SP-B incorporation into LNPs **(Fig. 1A)**.

LNP and Surf-LNPs were prepared at an N/P ratio of 6 and an aqueous-to-organic phase ratio of 1:1, and analyzed for their size, particle concentration, and zeta potential using Nanoparticle Tracking Analysis (NTA). Surf-LNPs demonstrated a slight yet statistically significant reduction in particle diameter compared to conventional LNPs (**Fig. 1B**). Both systems showed comparable zeta potentials (∼ -25 mV) (**Fig. 1C**). However, even at the same effective mRNA/lipid concentration, we observed a roughly order of magnitude increase in particle count of Surf-LNPs compared to conventional LNPs (**Fig. 1D**). The significant increase in particle count could be attributed to SP-B’s structural properties. The slight positive charge on SP-B (+5 – +8)^40^ could enable closer interaction with the mRNA cargo and faster, more efficient compaction than fully neutral helper lipid components. This could result in a more efficient, compact assembly of Surf-LNP particles, potentially yielding a higher particle count per unit mass of lipid components than conventional LNPs lacking SP-B. Both systems showed comparable and acceptable mRNA encapsulation efficiencies (**Fig. 1E**).

To assess particles’ stability in a physiologically relevant medium, Surf-LNPs or LNPs were prepared and incubated in simulated lung fluid (Gamble’s solution). While Surf-LNP remained at an order of magnitude higher total particle concentration, Surf-LNPs showed apparently lower colloidal stability with a 32.7 ± 3.3% reduction in particle count, opposed to 23.1± 10.4% in the case of LNPs after 4h of incubation (**Fig. S2**). To next consider the barrier function of mucus towards pulmonary LNP delivery, NP diffusion in secreted airway mucus was evaluated using fluorescence video microscopy and multiple-particle tracking, as described in our previous work (**Fig. 2A**)^42,43^. Airway mucus was obtained from air-liquid interface cultures of differentiated BCi-NS1 lung epithelial cells using previously established protocols. DiD-labeled LNPs or Surf-LNPs were added to mucus and imaged using confocal microscopy to evaluate their mobility in the mucus barrier. Representative trajectories show both LNP types were highly diffusive, presumably due to the DSPE-PEG shielding these particles from mucoadhesion and trapping in the mucus gel (**Fig. 2B**). From these trajectories, the mean squared displacement (MSD) for each particle was determined as MSD(τ)=〈(*x*^2^+*y*^2^)〉. Based on prior work illustrating how NP diffusion through mucus, as assessed by MPT, translates to *in vivo* biodistribution in the airways^44–46^, we considered particles highly mobile and likely to rapidly penetrate the mucus barrier if they achieved an MSD (τ=1 s) that is equal to or exceeds 1 µm^2^. While Surf-LNPs appeared to perform more consistently across measurements than LNP (**Fig. 2C**), both systems showed comparable mobility in mucus with the majority of individual particles (>50%) able to rapidly diffuse through mucus.

**Figure 2.**
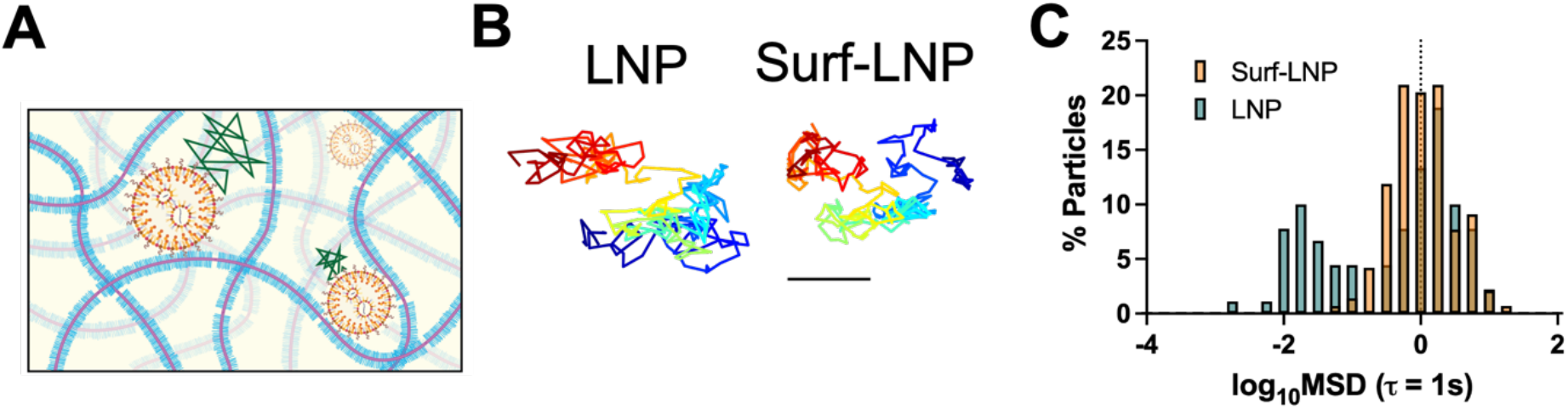
Diffusion of LNPs and Surf-LNPs in airway mucus. (**A**) Schematic of LNP diffusion through the mucus barrier. (**B**) Representative trajectories of convention LNP and Surf-LNP in mucus derived from human airway tissue cultures. Traces show 10 seconds of motion with a color scale to indicate time. The scale bar represents 1 µm. (**C**) Distribution of log based 10 of MSD at *τ* = 1 s (log_10_[MSD_1s_]) for LNP and Surf-LNP diffusion in mucus.

To compare the mRNA delivery efficiency of LNP and Surf-LNP, two cell lines were used: adenocarcinoma human alveolar epithelial cells (A549) and human airway basal cells (BCi-NS1). Each cell type was treated with either LNPs or Surf-LNPs at an mRNA dose of 1 µg/well for 24 h. In A549 cells, Surf-LNPs showed ∼8.5-fold increases in transfection efficiency compared to LNPs (**Fig. 3A**), as well as a ∼7-fold increase in protein expression (**Fig. 3B**). Both Surf-LNPs and LNPs demonstrated excellent cytocompatibility, with cell viability comparable to that of non-treated cells (**Fig. 3C**). **S**imilar trends were observed in BCi.NS1, where Surf-LNPs showed ∼6.5-fold increase in transfection efficiency (**Fig. 3D**), and 4 orders of magnitude increase in level of protein expression (**Fig. 3E**) compared to LNPs. Both particles demonstrated acceptable viability, with Surf-LNP showing slightly higher, though statistically non-significant cytocompatibility (**Fig. 3F**). The significant improvement in the ease of particle assembly, and hence a much higher particle count for Surf-LNPs at similar lipid and mRNA concentrations to conventional LNPs, may have contributed to the system’s enhanced in vitro transfection performance. Yet the previously reported endosomal escape-enhancing capabilities of poractant-α and its lipoprotein component, SP-B, may also have played a crucial role, which we attempted to assess in the next section.

**Figure 3.**
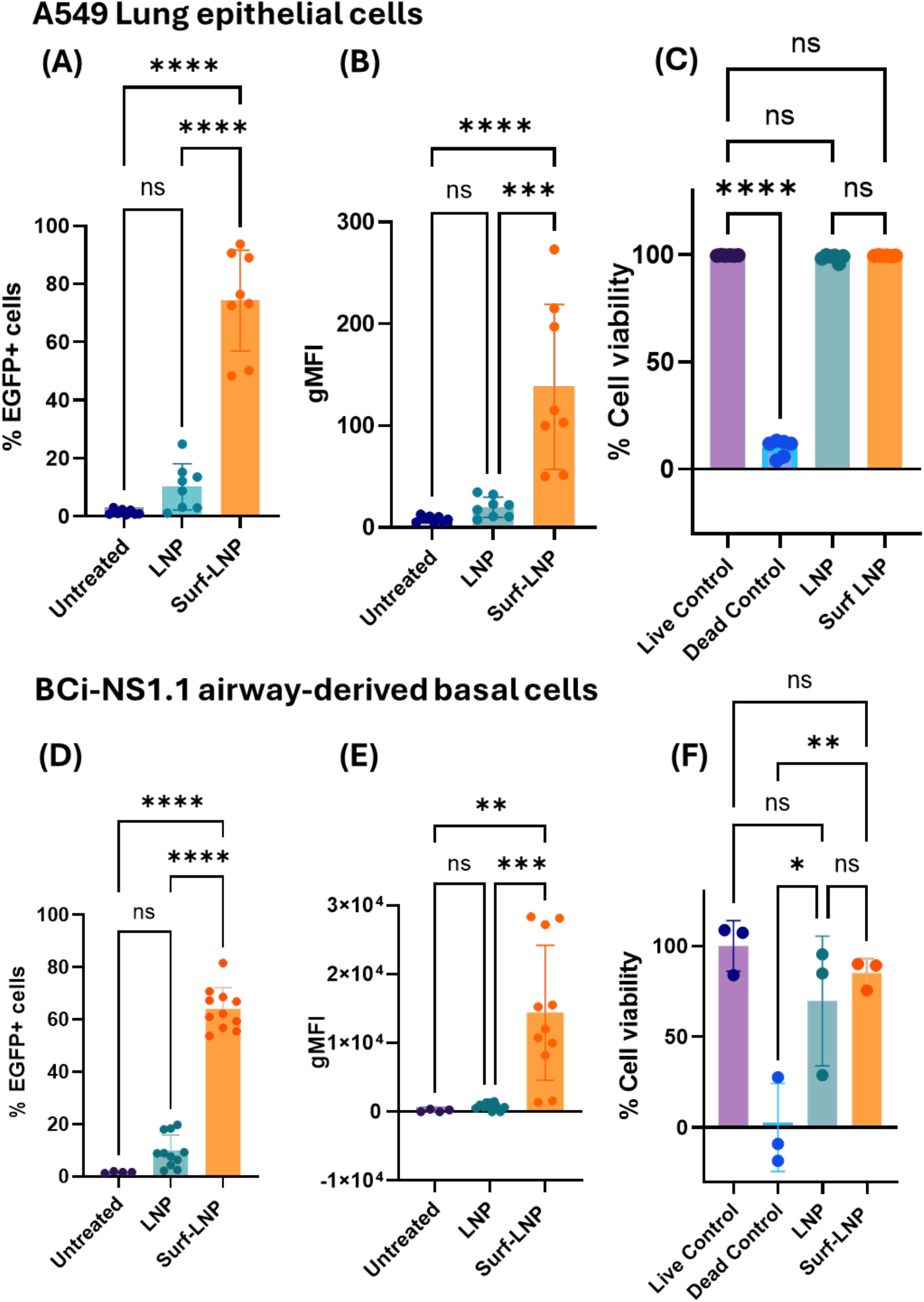
In vitro assessment of LNPs vs Surf-LNPs mRNA transfection. (**A**) Transfection efficiency (%EGFP+ cells) of mEGFP-loaded LNP / Surf-LNP in A549. (**B**) gMFI of A549 following treatment with mEGFP-loaded LNPs or Surf-LNPs. (**C**) Cytotoxicity of LNPs vs Surf-LNPs in A549. (**D**) Transfection efficiency of mEGFP-loaded LNP/ Surf-LNP in BCi-NS1.1. (**E**) gMFI of BCi-NS1.1 following treatment with mEGFP-loaded LNPs / Surf-LNPs. (**F**) cytotoxicity of LNPs vs Surf-LNPs in BCi-NS1.1. Data is displayed as Mean ± SD, (N=3), and analyzed by ANOVA where ns=non-significant, * p < 0.05, ** p < 0.01, *** p < 0.001, and **** p < 0.0001. gMFI; Geometric Mean fluorescence intensity, EGFP; Enhanced Green Fluorescent Protein.

The assessment of endosomal escape kinetics demonstrated higher as well as earlier uptake of DiD-labeled Surf-LNPs compared to plain LNPs (**Fig. 4A**). As early as 2h, prominent uptake of Surf-LNPs was observed; particles appeared uniformly distributed in the cytosol. However, the endosomal compartment proved to be tricky to stain at that time point. This could be due to early and efficient endosomal escape at this stage, leaving very few endosomal compartments intact. In contrast, DiD-labeled LNPs showed lower uptake at localized spots, and staining of endosomal compartments was feasible at 2h, presumably indicating slower, less efficient uptake and endosomal escape than with Surf-LNPs. At 6h, LNPs demonstrated a steady increase in uptake compared to their 2h point, yet they also demonstrated higher endosomal entrapment than Surf-LNPs (**Fig. 4B**).

**Figure 4.**
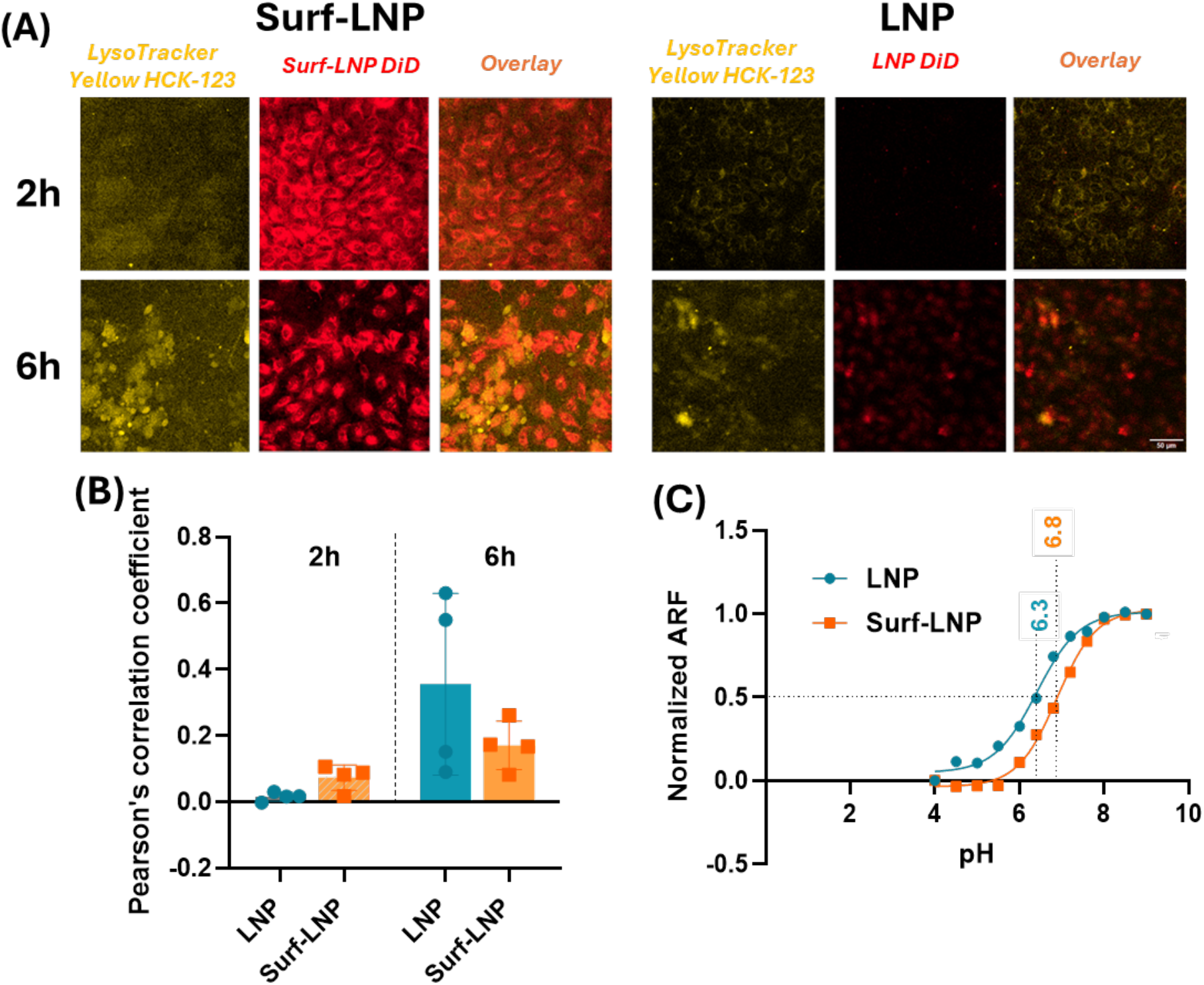
Colocalization analysis of LNP/Surf-LNP with endosomal compartments. (A) CLSM images demonstrating co-localization of DiD labelled LNP/Surf-LNP with Lysotracker Yellow HCK-123 staining endosomal compartments at 2h and 6h following LNP/Surf-LNP treatment in BCi-NS1 (Scale bar=50 µM) (B) Quantitative analysis of colocalization using Pearson’s correlation coefficient r=0 indicates no co-localization, r=1 indicates complete endosomal co-localization). (C) Apparent pKa of LNP vs Surf LNP. ARF; Average Relative Fluorescence.

LNPs have also demonstrated a lower pKa (6.3) than Surf-LNPs (6.8) (**Fig. 4C**), which is also consistent with the earlier endosomal escape observed for Surf-LNPs. We hypothesize that the higher apparent pKa of Surf-LNPs compared to LNPs could be attributed to the overall cationic charge of SP-B and SP-C in Proactant-α. Both lipoproteins could be stacking high up the lipid tails of anionic lipids and close to their anionic heads, involving these heads in subtle electrostatic interactions, which could render the ionizable lipids more susceptible to protonation at a higher pH than a lipid film where anionic lipids are more available to compete for the proton with the neighboring ionizable lipid. The earlier endosomal escape of Surf-LNP could thus be attributed to a range of mechanisms, in which the higher apparent pKa of Surf-LNP enables it to fuse with the endosomal lipid bilayer, thereby converting it to a hexagonal state more rapidly^47,48^. Simultaneously, as SP-B leaks from the particle, it can form channels in the endosomal membrane, further accelerating endosomal escape. Once the pH drops sufficiently in late endosomes, SP-B begins to attack the anionic components of the limiting membrane and ILVs, and escapes via either direct or back fusion.

As noted earlier, prior work identified SP-B can enhance gene delivery by acting as a potent endosomal escape enhancer. As an endosome matures, late endosomes become richer in anionic endogenous lipids such as bis(monoacylglycero)phosphate, phosphatidylserine, and phosphatidylinositol. Such anionic lipids are present both in the outer membrane of the late endosome and its ILVs present in late endosomes. Considering this, a dual mechanism was proposed in previous work by which SP-B-containing particles escape the endosome, both by virtue of SP-B’s cationic charge interacting with and disrupting the anionic lipids of either the outer membrane allowing the escape of the RNA cargo directly into the cytosol “direct fusion” or through fusion with the ILVs membranes, followed by “back fusion” with the limiting membrane and into the cytosol^32^. Biophysical evaluation has further demonstrated the essential role of SP-B in promoting the absorption of lung surfactant lipid components to the alveolar air-liquid interface as multilamellar films^49,50^. This function is facilitated by the fusogenic and surface-active properties of SP-B, which enable it to pack lipid films more closely together. Recent studies have also demonstrated that SP-B dimers tend to assemble into ring-like nanosized multimers through lateral migration in lipid films, especially in non-saturated lipids. Additionally, such rings can stack as channels that connect these lipid films in multilayers^51–53^. This mechanism could practically allow SP-B, once liberated from the Surf-LNP, to ‘punch nano-pores’ in the endosomal membrane that could facilitate mRNA escape. To this end, we plan to similarly evaluate these functions in LNP formulations with substitution of helper lipids with DSPC and inclusion of SP-B to compare to the Surf-LNP formulation in future studies.

To further evaluate pulmonary mRNA delivery *in vivo*, 8-week-old female BALB-c mice were intranasally inoculated with either DPBS (n=2), LNPs (n=5), or Surf-LNPs (n=5). Both LNP and Surf-LNP mice received an mCherry dose of 2.5 µg mRNA. We note this dose is lower than many previous studies on intranasal LNP delivery but was selected based on previous reports showing substantial inflammation at doses >0.5 µg mRNA following intranasal delivery^54^. After either 6 h or 24 h, the animals were sacrificed immediately and lung tissue was harvested for subsequent analysis. Based upon confocal imaging of lung tissue section, greater mCherry expression was observed in the tracheal lining following Surf-LNP administration as opposed to LNPs (**Fig. 5A, B**). A similar trend was observed in the lung parenchyma (**Fig. S4**), which demonstrated significantly higher expression of mCherry compared to LNP (**Fig. 5B**), though most expression was observed around bronchioles.

**Figure 5.**
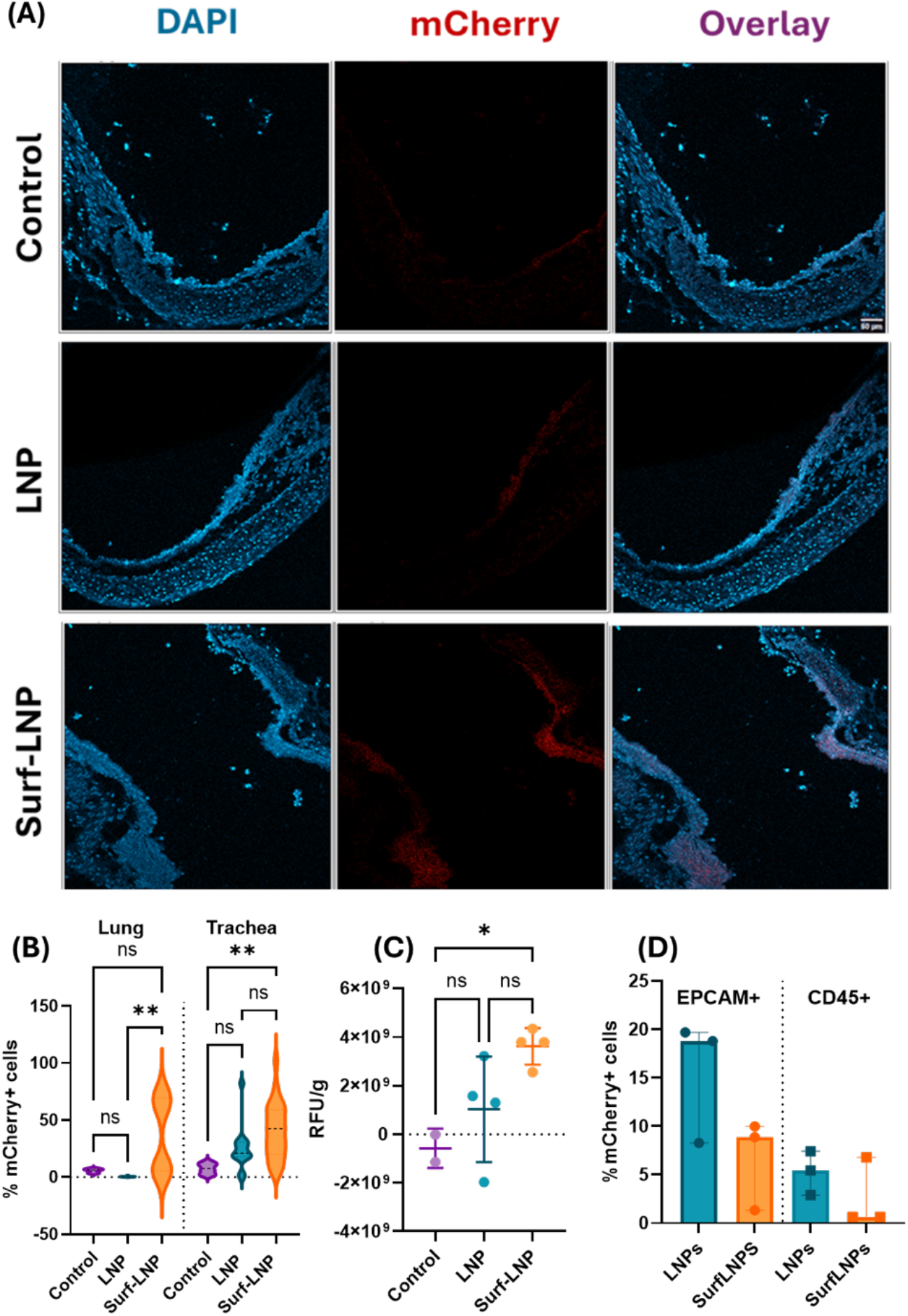
In vivo assessment of LNP and Surf-LNP mediated pulmonary mRNA delivery. **(A)** Fluorescent micrographs of the trachea from Balb-c mice 6h after treatment with mCherry expressing mRNA delivered using LNP and Surf-LNP (scale bar = 50 µm). **(B)** Quantification of %mCherry positive cells normalized to the total number of DAPI+ cell nuclei in fluorescent micrographs. **(C)** Bulk measurement of mCherry fluorescence in lung homogenates from untreated, LNP, and Surf-LNP treated mice. Data normalized to total weight of tissue. RFU = Relative Fluorescence Units. (**D)** Percentage of mCherry+ epithelial (EPCAM+) and immune (CD45+) cells as determined using flow cytometry. Data sets in B-D statistically analyzed by ANOVA: *p<0.05; **p<0.01; ns=not significant.

Tissue homogenates were also prepared for bulk assessment of mCherry expression and flow cytometry was performed to assess reporter expression in immune (CD45+) versus epithelial (EPCAM+) cell populations. Whole lung homogenates from tissues collected 24h post-treatment showed a significant increase in mCherry expression in Surf-LNP treated groups compared to untreated controls (**Fig. 5B**). LNP showed a more variable expression between replicates and no significant difference in the level of mCherry expression with either PBS or Surf-LNP-treated groups. Interestingly, flow cytometric analysis of mCherry expression in EPCAM+ vs CD45+ cells in a single cell suspension of the lung tissue showed that LNP transfected both EPCAM+ and CD45+cells, yet Surf-LNP appeared to predominantly transfect EPCAM+ cells (**Fig. 5C**). More data is still required to fully elucidate this finding, yet several accounts in the literature have demonstrated the propencity of smaller nanoparticles <100-200 nm to be taken up by endocytosis by non-immunogenic cell populations (such as epithelial cells), as opposed to larger (>200 nm) particles more commonly taken up phagocytosis by CD45+ macrophages or other antigen presenting cells^55–57^. Inflammatory cytokine response assessment in the supernatant of lung homogenates after 6h of particle treatment demonstrated that neither LNPs nor Surf-LNPs induced a significant increase in inflammatory cytokines (TNF-α, IL-1ß, IL-6) compared to the PBS-inoculated group (**Fig. S5**).

In conclusion, we demonstrated that replacing the standard helper lipid (DSPC) of commercial LNPs with a biomimetic helper lipid cocktail of commercially available lung surfactant (poractant-α) resulted in a logarithmic increase in particle count due to improved particle assembly. This improvement could be attributed to the unique saposin-like structure of SP-B, which may have promoted a more compact and efficient packing of the surfactant-containing Surf-LNP compared to conventional LNP. Furthermore, the resulting Surf-LNP showed earlier and higher endosomal escape, potentially due to the superior fusogenic properties of SP-B, as well as the elevated apparent pKa of Surf-LNPs compared to LNPs. Due to enhancement in endosomal escape, Surf-LNPs significantly outperformed conventional LNPs in both transfection efficiency and protein expression levels. Our findings highlight that incorporating lung-derived helper and structural LNP components, which actively facilitate endosomal escape, could enhance LNP performance. We also observed a ∼2-fold improvement in mRNA expression *in vivo* following intranasal delivery of Surf-LNPs as compared to LNPs and a transfection profile that notably favored epithelial over immune cells. As such, Surf-LNPs warrants further investigation for protein replacement therapy applications which would benefit from targeting epithelial cell subsets. Given this approach could be used in a wide variety of newly emerging LNP formulations with greater potency, we plan to adapt this formulation approach in combination with other ionizable lipid platforms with the aim of further improving pulmonary mRNA therapeutics for chronic lung diseases.

## ASSOCIATED CONTENT

**Supporting Information** includes Materials & Methods as well as supporting data in Figures S1-S5:

**Figure S3**. (A) Comparative compositions of physiological human lung surfactant (ref 30) vs commercially available porcine lung surfactant cocktail Poractant-α (ref. 29). (B) Chemical structures of DSPC (helper lipid in commercially available LNPs) and DPPC (main phospholipid component in Poractant-α. (C) Diagrammatic illustration of LNP vs Surf-LNP composition. PC: phosphatidyl choline, PL: phospholipids, PG: phosphatidyl glycerol.

**Figure S4**. Particle stability demonstrated as a function of changes in particle count over time following 4h incubation in simulated lung fluid (Gamble’s solution).

**Figure S5**. In vitro assessment of LNPs vs Surf-LNPs in A549 cells. Confocal images demonstrating GFP expression in untreated, LNP-treated, and Surf-LNP-treated A549 (scale bar= 50 µm).

**Figure S4**. Representative confocal images of lung parenchyma of Balb-c mice after 24h of treatment, respectively (scale bar = 50 µm)

**Figure S5**. Cytokine levels in lung tissue homogenates from LNP and Surf-LNP treated mice measured by ELISA.

## AUTHOR CONTRIBUTIONS

The manuscript was written through contributions of all authors. All authors have given approval to the final version of the manuscript.

## CONFLICTS OF INTEREST

S. N. and G. A. D. have a pending patent based on the lipid nanoparticle formulation described in this manuscript.

## FUNDING SOURCES

This project was funded by the National Institutes of Health (R01 HL160540).

## ACKNOWLEDGMENT

We acknowledge the BioWorkshop core facility and Dr. Christian Pick in the Fischell Department of Bioengineering at the University of Maryland for use of their microplate reader and flow cytometer.

