## Supplement for "Lung surfactants as a component of lipid nanoparticles for pulmonary mRNA delivery"

**Supporting Information**

### MATERIALS & METHODS

*Nanoparticle formulation*

LNPs/Surf-LNPs were prepared as follows: an ethanolic solution of Dlin-MC3-DMA (MedChemExpress, NJ, USA), Cholesterol (Sigma Aldrich, USA), DSPC (Avanti Polar Lipids, AL, USA) (for LNPs) or Proactant-α (for surf LNPs, calculated based on DPPC content)(Creative Biolabs, NY, USA), DSPE-PEG2000K (Avanti Polar Lipids, AL, USA) was prepared at molar ratios of 50%, 38.5%,10%,1.5%, respectively. Messenger RNA (mEGFP for in-vitro assays, mCherry for in-vivo assays) (Trilink, CA, USA) was dispensed in an acetate buffer (100mM, pH=5). The ethanolic and aqueous phases were mixed by pipetting at an N/P ratio of 6, and an aqueous to organic phase ratio of 3:1 v/v. Immediately after, particle suspension was diluted 10X using Dulbecco's Phosphate Buffer Saline (DPBS, pH=7.4). The diluted suspension was then washed twice with centrifugal filtration using Amikon 100k and re-diluted to the original volume using DPBS. For multiple particle tracking experiments, DiD-labeled particles were prepared similarly except that a 0.3% molar ratio of DiD (Sigma Aldrich,USA) was included in the lipid mixture. That contribution was deduced from the molar contribution of cholesterol in the non-labeled particle variety.

*Colloidal properties*

Particles were analysed for their size, count, and zeta potential using Nanoparticle Tracking Analysis (NTA)(ZetaView®, Particle Metrix, NC, USA). Briefly, either LNPs or Surf-LNPs were diluted 1000-fold in DPBS, and videos of the particles were acquired at 25 °C at 11 different focal positions within the sample.

*Particle stability in simulated lung fluid (Gamble's solution)*

Surf-LNPs or LNPs prepared as previously described were incubated in Gamble's solution (Pickering Laboratories Inc., CA, USA) at a ratio of 1:10 v/v, at 37 °C with shaking at 70 RPM for 30 minutes. Following this, the incubation solution was diluted 300-fold in DPBS to quench the reaction. Particle count was then assessed using NTA as previously described.

*Encapsulation efficiency*

LNPs and Surf-LNPs were analyzed for their mRNA encapsulation efficiency using the Ribogreen assay (ThermoFischer Scientific, USA). Either particle was prepared as previously described. The filtrate passing through the Amikon was collected and analyzed for unencapsulated mRNA content using the Ribogreen assay according to the manufacturer's protocol. The amount of free mRNA interacting with Ribogreen dye was assessed spectrophotometrically at $\lambda_{ex}$=500nm, $\lambda_{em}$=525 nm using Multimode Microplate Reader (Tecan, Männedorf, Switzerland). Then the encapsulation efficiency (EE%) was calculated according as EE% = [(Total mRNA – Unencapsulated mRNA)/(Total mRNA)] x 100%.

*Apparent pKa*

Apparent pKa was assessed for LNPs or Surf-LNPs using the Lipid Launch Apparent pKa kit (Cayman Chemical, MI, USA). Briefly, Surf-LNP or LNP were prepared as previously described. Samples were washed twice using a 100 kDa MWCO Amikon centrifugal filter (Millipore Sigma, USA) with DPBS at

2150 g for 5 min. Particles were brought to a final concentration of 1.3 µg mRNA/µL. Surf-LNPs were then diluted 20X, whereas LNPs were used undiluted. In 96-well plates, particle suspensions were mixed with the kit-provided buffers at an 18:1 v/v ratio, with the following pH gradient: 4, 4.5, 5, 5.5, 6, 6.4, 6.8, 7.2, 7.6, 8, 8.5, and 9. Following this, the TNS probe solution was added to the reaction mixture at a 1:20 v/v ratio, and the reaction was allowed to proceed for 20 minutes on an orbital shaker at 60 RPM. The fluorescent signal of the samples was assessed using a plate reader ($\lambda_{ex}$=320 nm, $\lambda_{em}$ = 450 nm). Average relative fluorescence (ARF) is determined by subtracting the average fluorescence of the pH 9.0 wells from all other wells of a sample. ARF is then plotted against pH, and pKa is calculated at the pH corresponding to 50% maximal fluorescence.

*Multiple particle tracking analysis of LNP diffusion through mucus*

To generate mucus for these studies, BCi-NS1.1 human airway epithelial (HAE) cells were first expanded on tissue culture plastic in Pneumacult-Ex or Pneumacult-Ex Plus medium (no. 05008 or 05040, StemCell Technologies). BCi-NS1.1 HAE cells were then seeded ($3.3 \times 10^4$ cells per well) on rat tail collagen type 1-coated permeable Transwell membrane supports (6.5 mm; no. 3470, Corning, Inc.) and differentiated in Pneumacult-ALI medium (no. 05001, StemCell Technologies) at an air–liquid interface (ALI) for at least 28 days until matured into a pseudostratified mucus-producing and ciliated epithelium. To collect the apically secreted mucus, HAE cultures were washed with phosphate buffered saline (PBS) to remove the accumulated mucus from the apical surface for collection. The collected mucus was filtered using Amicon ultra centrifugal filter units with a 100 kDa cutoff to remove excess PBS. The resulting mucus was stored at -80 °C until time of use. To evaluate the movement of LNPs in airway mucus, 1 µL of each type of particle was added to 20 µL of airway mucus and placed on a slide in the middle of a vacuum grease-coated O-ring. Slides were equilibrated for 30 minutes at room temperature prior to fluorescence imaging with a Zeiss Confocal LSM 800 microscope equipped with a 63x water-immersion objective. Multiple 10-second videos were recorded at 33.3 frames per second for each sample. Fluorescence microscopy video files were processed using a previously developed MATLAB code capable of tracking multiple particles and calculating the MSD. The MSD was calculated as $\langle MSD(\tau) = (x^2 + y^2) \rangle$, for each particle.

*In-vitro Transfection*

Adenocarcinoma human alveolar basal epithelial cells (A549) and an immortalized human airway basal cell line (BCi-NS1) were used for our studies. A549 cells were passaged, grown, and maintained in F12K medium supplemented with 10% FBS and 1% Pen/Strep. Bci-NS1 were passaged, grown, and maintained in EX-plus medium supplemented with PneumaCult™-Ex Plus 50X Supplement, 96 µg/L hydrocortisone (Stem Cell Technologies, MA, USA), 1% Pen/Strep, 0.5%Amphotericin (Sigma Aldrich, USA). Cells were consistently kept at 37 °C and 5% $CO_2$. In either cell line, the transfection procedure was the same. Cells were seeded at a density of 25000 cells/well in 24-well plates. They were allowed to grow to 70-80% confluenec over the course of 48h in their respective growth medium. Cells were then treated with either LNPs, Surf-LNPs, or JetMessenger (Polyplus, NY, USA) with mRNA doses of 1 µg/well in OptiMEM for 24h. Cells incubated in OptiMEM (ThermoFischer Scientific, USA) without treatment served as the untreated control. Afterward, OptiMEM, with or without treatment, is removed; cells are washed twice with DPBS and further incubated in their standard growth medium for another 24h. Eventually, cells were detached with Trypsin-EDTA (Sigma Aldrich, USA) and analyzed for mEGFP expression using a FACS Celesta (BD Biosciences, CA, USA).

*Endosomal Escape Kinetics*

BCis-NS1 were used for endosomal escape kinetic studies. Cells were seeded as previously described for in vitro transfection trials, except that they were seeded on top of microscopic cover slides in the 24-well plates. Cells were then treated with either DiD-labeled LNPs or Surf-LNPs in OptiMEM (1µg mRNA/well). At 2h, 6h, or 24h, treatments were removed, and cells were stained with 80 nM Lysotracker Yellow HCK-123 (ThermoFischer Scientific, USA) in the previously described BCi-NS1 growth medium for 1h at 37 °C, 5% CO2. Following this, the respective coverslides were retrieved, mounted on microscopic slides using Vectashield (Vector Labs, CA, USA), and visualized using Confocal Laser Scanning Microscopy (Zeiss LSM 780, Zeiss, USA). The acquired images were analyzed for DiD (particle) and Lysotracker (endolysosome) colocalization over time using the Colocalization Threshold plugin on ImageJ.

*Intranasal Delivery of LNPs*

Twelve 8-week-old female BALB-c mice were intranasally inoculated with either DPBS (n=2), LNPs (n=5), or Surf-LNPs (n=5). Both LNP and Surf-LNP mice received an mCherry dose of 2.5 μg mRNA per animal in a total volume of 20 μL (10 μL/nostril). Doses were administered using a 10 μL pipette to the animal under anesthesia. After 24 h, live animals were imaged using an IVIS Spectrum In Vivo Imaging System (Revvity). After either 6h or 24h, animals were immediately sacrificed. Afterward, lungs (at 24h)and trachea (at 6h) were harvested. For quantitative assessment of mCherry expression, lungs were homogenized in T-PER lysis buffer (ThermoFischer Scientific, USA) supplemented with protease inhibitor cocktail (ThermoFischer Scientific, USA). Briefly, lungs were mixed with the buffer at 5% w/v and diced using a Pestle Motor Mixer. Tissue homogenates were then centrifuged at 10000 g for 5 mins. Supernatant was spectrophotometrically assessed for mCherry fluorescence at $\lambda_{ex}$=587 nm, $\lambda_{em}$=610 nm using Multimode Microplate Reader (Tecan, Männedorf, Switzerland).

Flow cytometric analysis of mCherry expression in immune vs epithelial cell populations was performed. Briefly, Lung tissue was finely diced, then incubated in a solution of collagenase IV (0.5 mg/mL) (Sigma-Aldrich, USA) and Dnase I (10 μg/mL) (Sigma-Aldrich,USA) in RPMI at 37 °C, 5% CO2, shaking at 50 RPM for 45 minutes. Tissue digest was then passed through a 70 μm cell strainer (Millipore Sigma, USA). Cells were then collected by centrifugation at 300 g for 10 minutes. The pellet was washed with PBS, then treated with RBC lysis buffer for 3 minutes on ice. The reaction was halted by 4x dilution in cold PBS, followed by washing.

Cells were stained for either CD45 or EpCAM as follows. First, cells were incubated with CD16/CD32 Monoclonal antibody for 10 minutes on ice to perform Fc blocking. Cells were then washed with PBS and incubated with either FITC-labeled anti-CD45 or anti-EpCAM (Invitrogen, Thermo Fisher Scientific, USA) for 45 minutes on ice. Cells were washed and permeabilized for intracellular staining using Cytoperm/Wash for 15 minutes. Cells were then incubated with Alexa Fluor 647-labeled anti-mCherry monoclonal antibody (Invitrogen, Thermo Fisher Scientific, USA) for 30 minutes on ice. Cells were then analyzed using FACS Celesta (BD Biosciences, CA, USA). For confocal imaging, lungs or tracheas from each treatment group were mounted in OCT, sectioned on a cryomicrotome, and mounted on microscopic slides. The mounted tissue was fixed with 4% PFA, washed with PBS, permeabilized with 0.5% Triton X-100 (v/v) in PBS, and then stained with 300 nM DAPI (Sigma Aldrich, USA) in PBS. The tissues were then imaged with CLSM to visualize mCherry expression. % mCherry-expressing cells, relative to the total number of cells per image, was calculated from CLSM images using ImageJ, with the untreated control used to set the threshold for mCherry expression.

SUPPLEMENTARY FIGURES

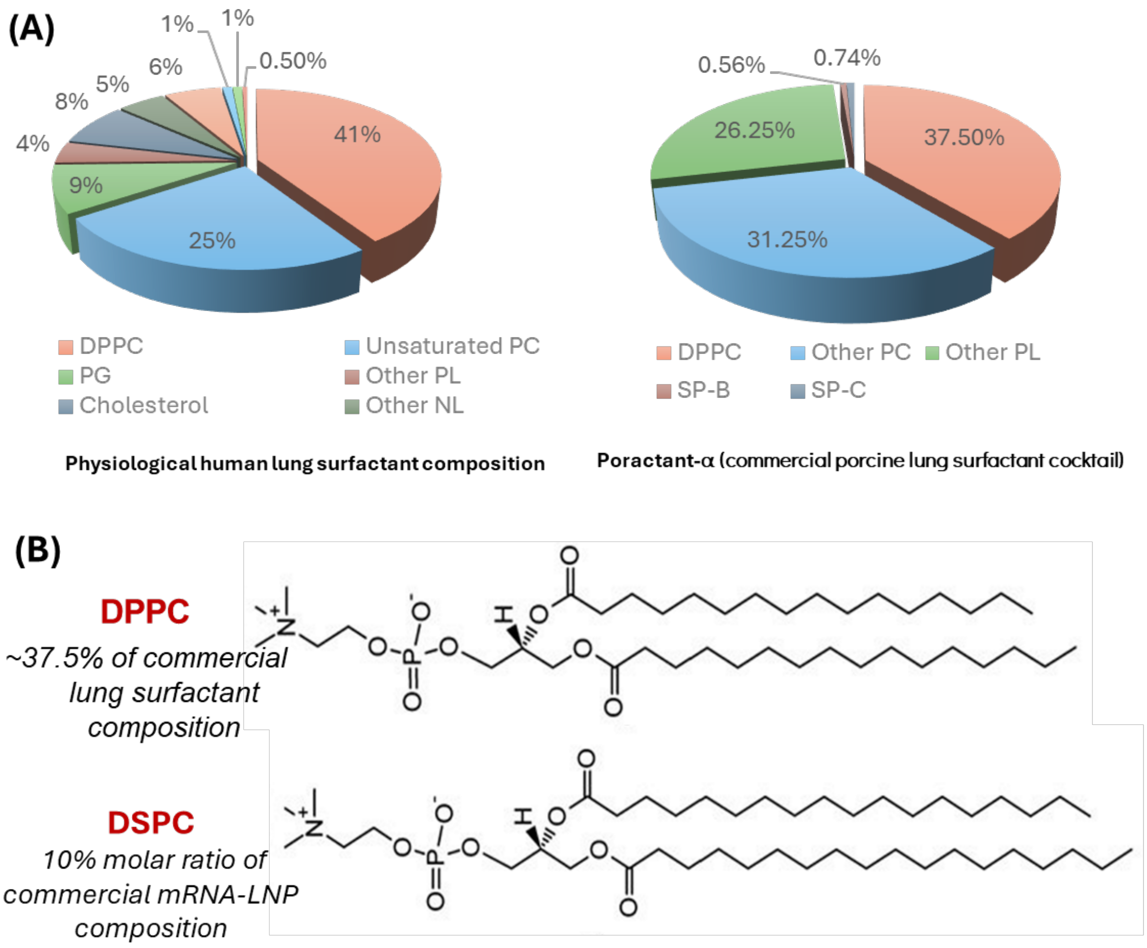


**Figure S1.** (A) Comparative compositions of physiological human lung surfactant (ref 30) vs commercially available porcine lung surfactant cocktail Poractant-α (ref. 29). (B) Chemical structures of DSPC (helper lipid in commercially available LNPs) and DPPC (main phospholipid component in Poractant-α. (C) Diagrammatic illustration of LNP vs Surf-LNP composition. PC: phosphatidyl choline, PL: phospholipids, PG: phosphatidyl glycerol.

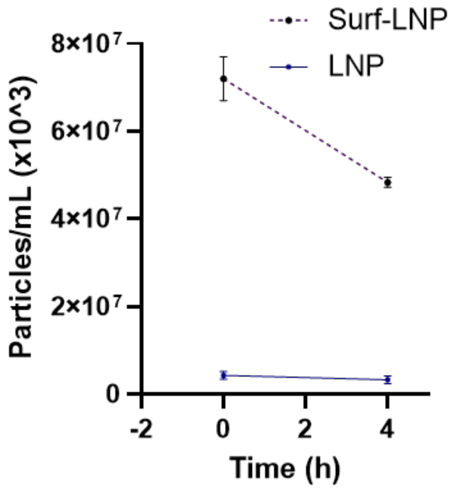


**Figure S2.** Particle stability demonstrated as a function of changes in particle count over time following 4h incubation in simulated lung fluid (Gamble's solution).

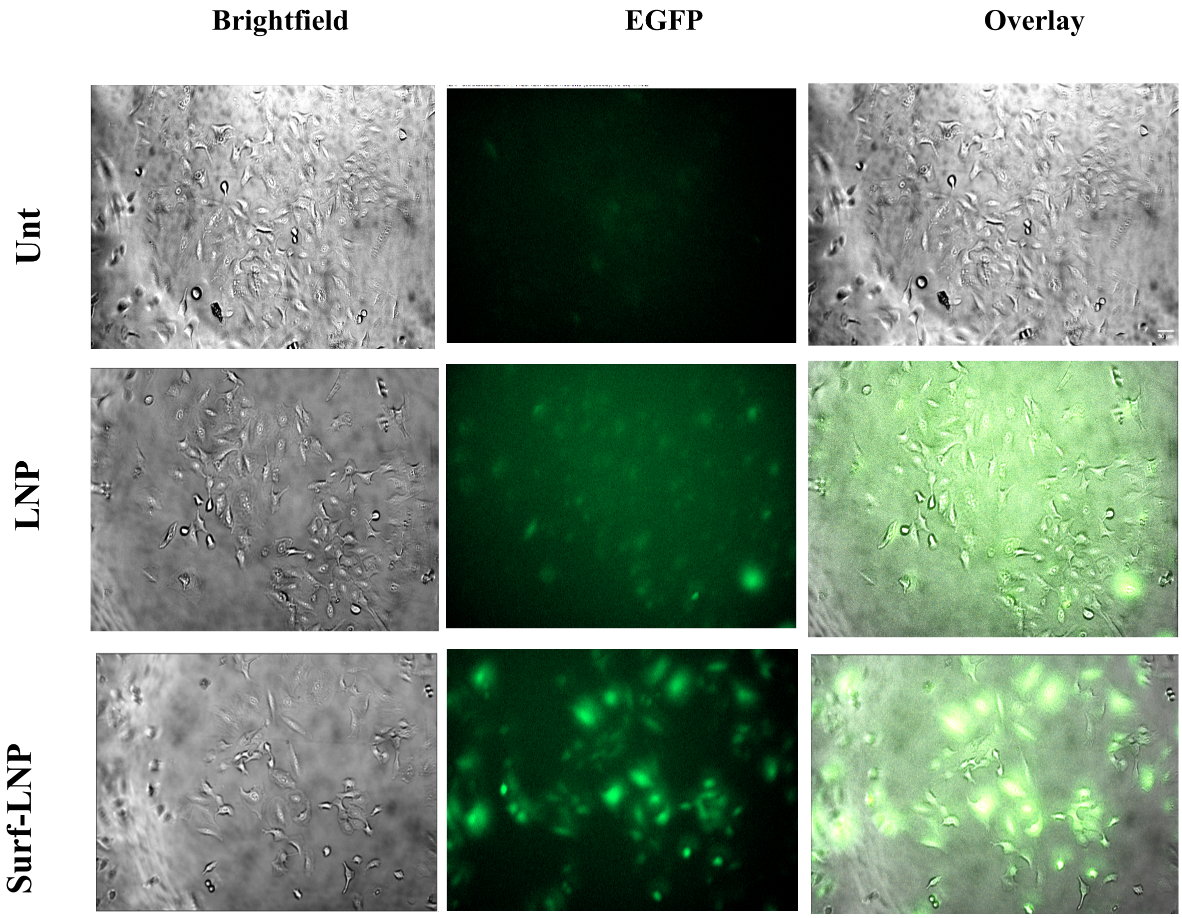


**Figure S3.** In vitro assessment of LNPs vs Surf-LNPs in A549 cells. Confocal images demonstrating GFP expression in untreated, LNP-treated, and Surf-LNP-treated A549 (scale bar= 50 μm).

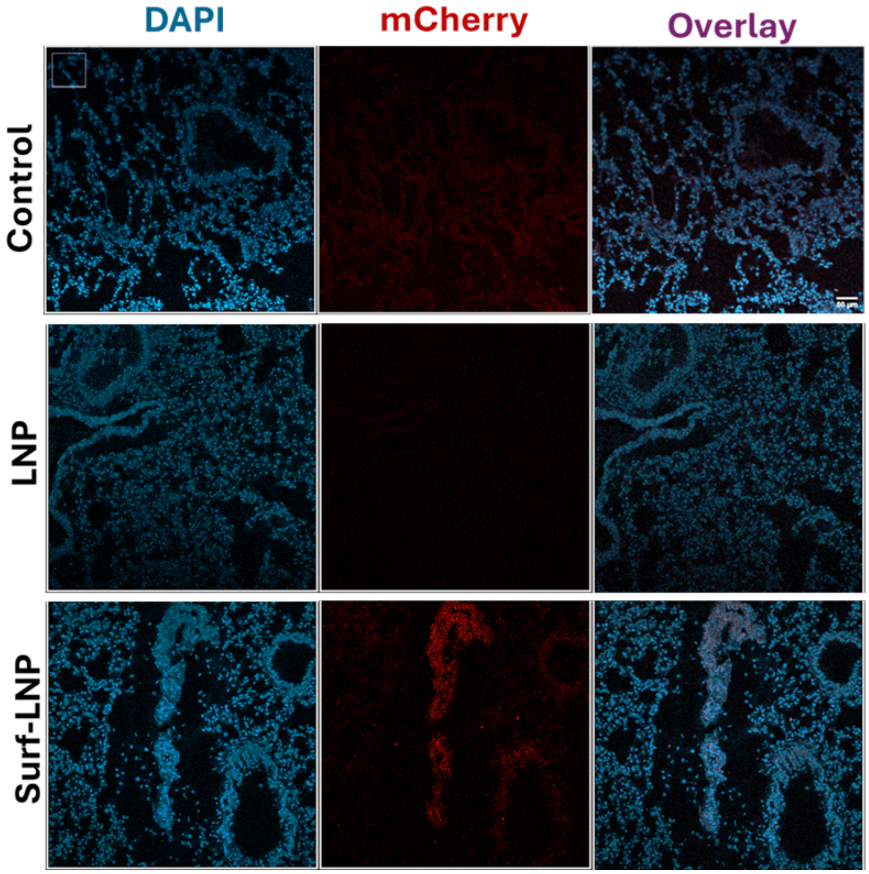


**Figure S4.** Representative confocal images of lung parenchyma of Balb-c mice after 24h of treatment, respectively (scale bar = 50 µm)

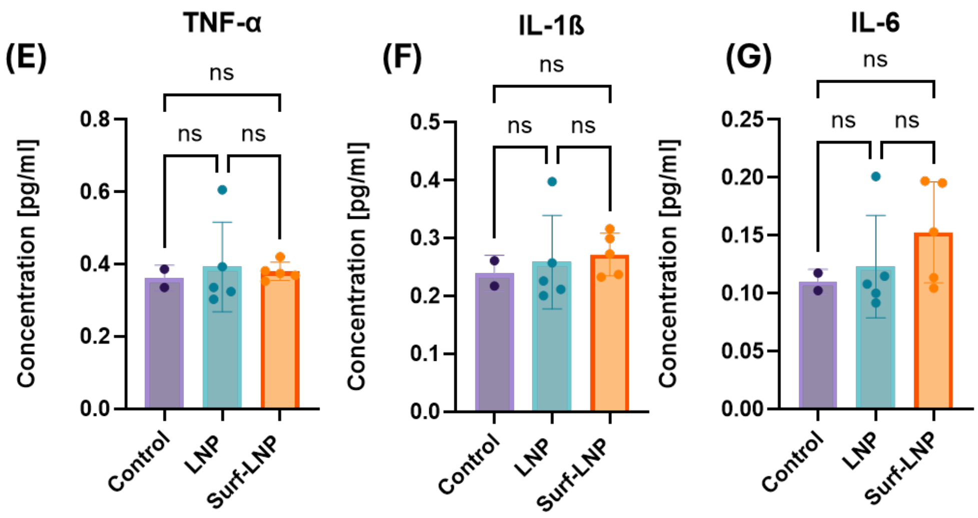


**Figure S5.** Cytokine levels in lung tissue homogenates from LNP and Surf-LNP treated mice measured by ELISA. Datasets statistically analyzed by ANOVA: ns=not significant.
